# Implications of back-and-forth motion and powerful propulsion for spirochetal invasion

**DOI:** 10.1101/2020.04.27.065318

**Authors:** Keigo Abe, Toshiki Kuribayashi, Kyosuke Takabe, Shuichi Nakamura

## Abstract

The spirochete *Leptospira* spp. can move in liquid and on a solid surface using two periplasmic flagella (PFs), and its motility is an essential virulence factor for the pathogenic species. Mammals are infected with the spirochete through the wounded dermis, which implies the importance of behaviors on the boundary with such viscoelastic milieu; however, the leptospiral pathogenicity involving motility remains unclear. We used a glass chamber containing a gel area adjoining the leptospiral suspension to resemble host dermis exposed to contaminated water and analyzed the motility of individual cells at the liquid-gel border. Insertion of one end of the cell body to the gel increased switching of the swimming direction. Moreover, the swimming force of *Leptospira* was also measured by trapping single cells using an optical tweezer. It was found that they can generate ∼17 pN of force, which is ∼30 times of the swimming force of *Escherichia coli*. The force-speed relationship suggested the load-dependent force enhancement and showed that the power (the work per unit time) for the propulsion is ∼3.1×10^−16^ W, which is two-order of magnitudes larger than the propulsive power of *E. coli*. The powerful and efficient propulsion of *Leptospira* using back-and-forth movements could facilitate their invasion.

## Introduction

Motility is a crucial virulence factor for pathogenic bacteria^1^. For example, a motility-deficient mutant of *Vibrio cholerae* is attenuated due to the decreased invasion efficiency of the epithelium^2^. In some flagellated bacteria, not only motility but also the flagella themselves are essential as an adhesin, and *Salmonella* Enteritidis attaches to the host tissue via peritrichous flagella, resulting in colonization and clinical outcomes^3^. Although spirochetes, such as *Borrelia burgdorferi* (the Lyme disease)^4^ and *Brachyspira hyodysenteriae* (swine dysentery)^5^, also utilize motility during infection, their flagella exist beneath the outer membrane known as the periplasmic flagella (PFs), and the spirochetal flagella are not involved in pathogenicity directly. Instead, improvement of swimming ability^6^ and diverse adherence^7^ in viscoelastic environments are believed to be responsible for their colonization and dissemination within hosts.

The genus *Leptospira* is a member of spirochetes, and the pathogenic species cause a worldwide zoonosis leptospirosis. Pathogenic *Leptospira* cells are maintained in the proximal renal tubules of reservoir rodents. When the hosts urinate, they spread the spirochetes into environments; many mammals, including humans, are percutaneously infected by contact with the contaminated soil and water^8,9^. *Leptospira* spp. have a right-handed spiral cell body and exhibit curvatures at both ends (Fig. 1A). The spirochete can swim in liquid and crawl on surfaces using two PFs (one PF/cell end) (Fig. 1B). The morphology of the cell ends frequently change between a spiral and hook shape: an asymmetric configuration of spiral and hook shapes at the anterior and posterior cell ends, respectively, propels the cell unidirectionally (Fig. 1B)^10–13^. As with other motile species, the motility of *Leptospir*a spp. is closely related to their pathogenicity^14,15^, although how it contributes as a virulence factor in the spirochete remains to be elucidated.

**Fig. 1.**
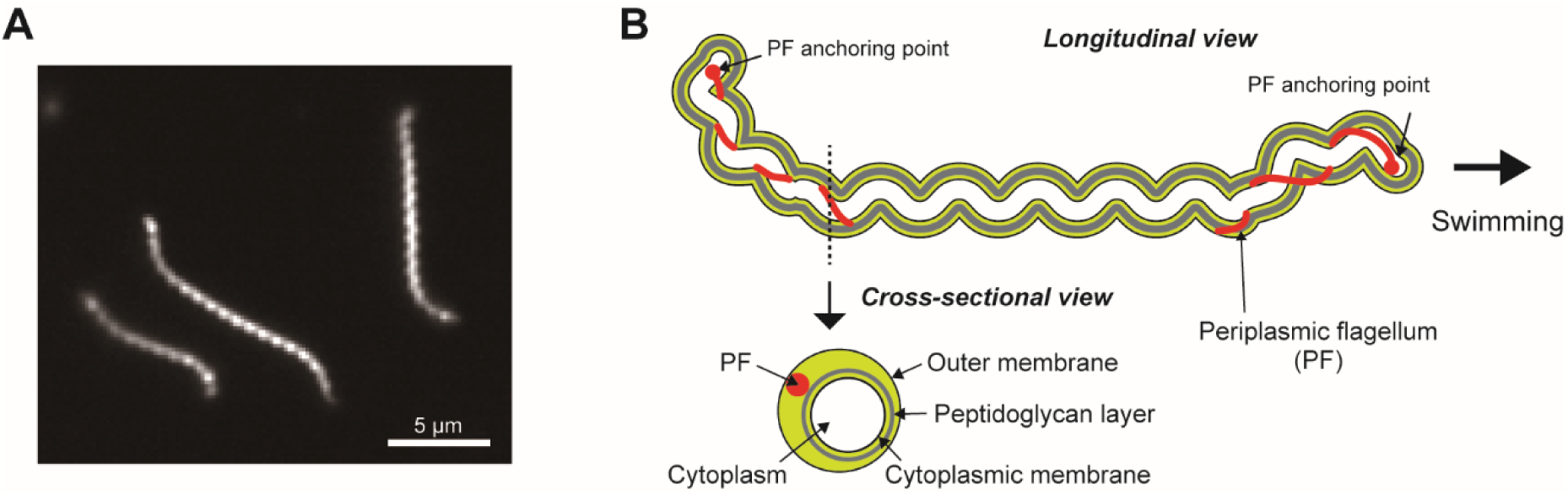
Morphology of *Leptospira*. (A) Dark-field micrograph of *Leptospira kobayashii*. (B) Schematic diagram of the cell structure of *Leptospira* spp.; a swimming cell exhibiting the spiral anterior end (right side of the diagram) and hook-shaped posterior end (left side) is illustrated.

As a possible explanation for the motility dependence in the pathogenicity of *Leptospira*, we recently showed that the spirochete increased the frequency of changing the swimming direction with the elevated viscosity of polymer solutions, and the resultant limitation of their net migrations could be involved in the accumulation of the spirochete in epithelial mucosa *in vivo*^12^. In this study, hypothesizing that the phenomenon observed in polymer solution for percutaneous infection of *Leptospira* is significant, we investigated their behaviors at the border of liquid and gel phases to mimic the skin dermis being exposed to contaminated environmental water. Furthermore, we expected there would have to be a sizeable propulsive force for the invasion and, thus, performed force measurements of swimming spirochetes using an optical tweezer. Our experiments showed that there is an enhancement of swimming reversal when only one end of the cell body is inserted into the gel, and much larger swimming force of *Leptospira* than the known values of exoflagellated bacteria. The results suggest that powerful and efficient swimming with repeated trial-and-error allows *Leptospira* to obtain a smooth passage for penetration through the host dermis.

## Results

### Swimming reversal

We observed the spirochetes using a flow chamber containing liquid medium and agar that were adjoining, in order to investigate the behaviors of *Leptospira* during penetration of viscoelastic environments (Fig. 2A). *Leptospira* cells show relatively smooth swimming in liquid (Figs. 2B left, and 2C left). In contrast, when one end of the cell body is inserted into agar, they frequently changed direction (Figs. 2B right, and 2C right). The significant difference in reversal frequencies between cells in liquid and at the liquid-agar border is shown in Fig. 2D. Video 1 shows that a *Leptospira* cell succeeds in penetrating agar after several back-and-forth movements. Enhancement of swimming reversal at the liquid-agar border was also observed in different species of *Leptospira* (Video 2 for *L. interrogans* and Video 3 for *L. biflexa*), suggesting that the phenomenon is shared among the genus. For pathogenic species, exploring more accessible routes in the dermis, such as structurally disturbed parts due to injury, using such a “trial and error” method could be involved in percutaneous invasion (Fig. 2E).

**Fig. 2.**
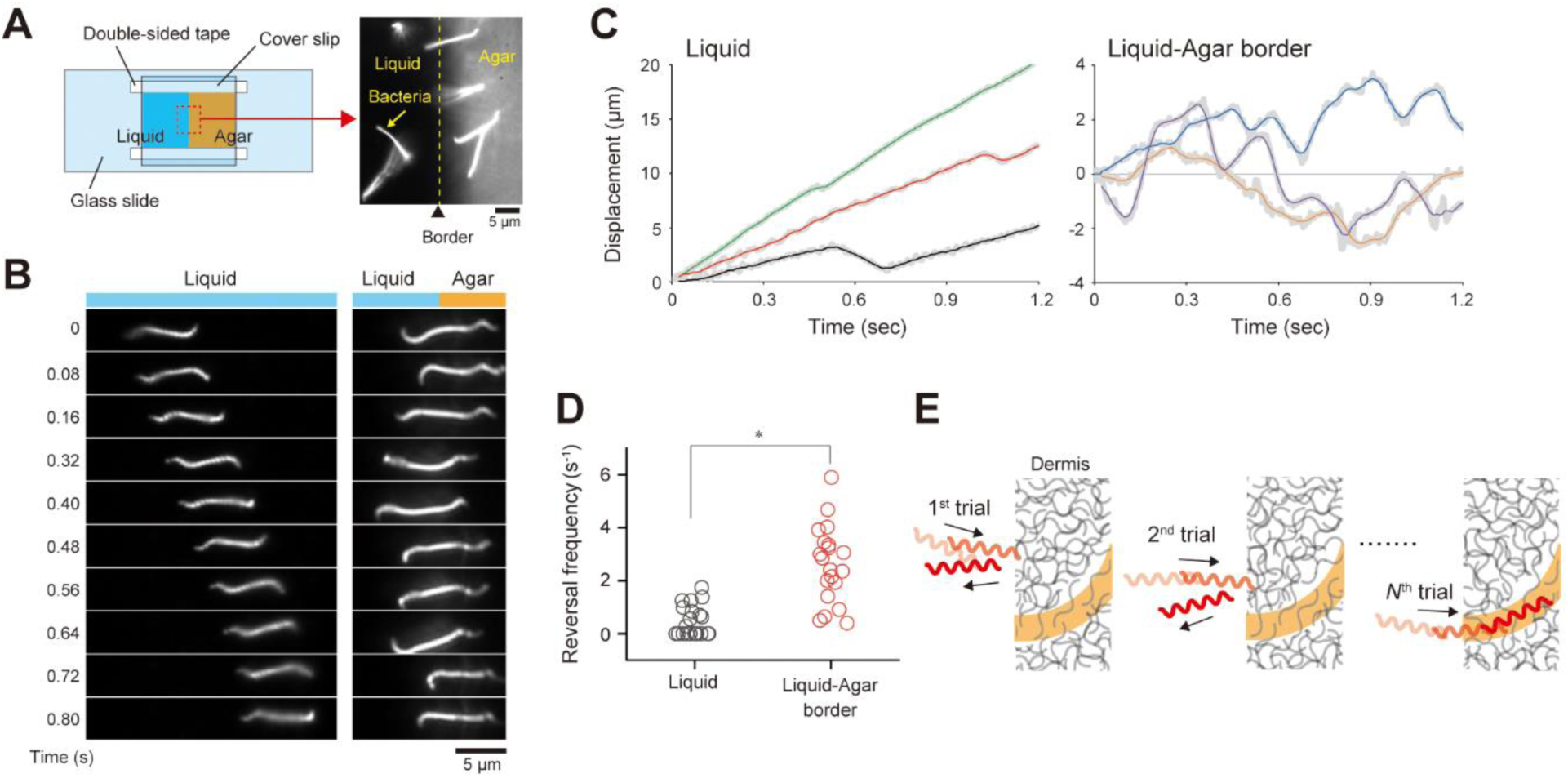
Swimming reversal. (A) Illustration of a flow chamber in which liquid and agar areas are contiguous. The microscopic image shows the liquid-agar border. Bacteria swam in liquid at the beginning of the experiment and then penetrated the agar area. (B) Time courses of the cell-movements observed in liquid (left) and at the liquid-agar border (right). (C) Displacements of individual cells. Example data obtained from three *L. kobayashii* cells are shown, each in liquid and at the border. Thick gray lines indicate raw data obtained by determining cell positions at 4 ms intervals, and thin colored lines are the results of a 12-data-point moving average. (D) Comparison of the reversal frequency determined from the cell displacement data. The results of liquid highlight no reversals occurred during observation. Mann-Whitney U test showed significant difference (\**P* < 0.05); n = 24 cells for liquid, and n = 20 cells for the border. (E) A plausible contribution of frequent swimming reversals for invasion. Back-and-forth movements give the cell chance to find a more accessible route in heterogeneous dermis structure (orange). Video 1 demonstrates “trial-and-error” at the liquid-agar border by swimming reversals.

### Force measurement for the *Leptospira* swimming

We trapped a microbead attached to swimming cells with an optical tweezer for measurement of the swimming force of *Leptospira* (Figs 3A-B). The bead displacements were converted to the swimming force (*F*) by considering the balance with the trapping force (*F*_*trap*_) and the drag force (*F*_*drag*_) exerted on the bead, *F* = *F*_*trap*_ + *F*_*drag*_ (Figs. 3C-D, and Video 4). The swimming force increased with the cell displacement and reached saturation when the cell was stalled by the restoring force of the laser trap. The stall force widely differed among the measured cells, and the averaged force-time curve showed a stall force of 16.6 ± 2.2 pN (mean ± standard error; n = 24 cells) (Fig. 4A). A model experiment in which a tungsten coil, mimicking the shape of *Leptospira*, was rotated in a rotational magnetic field, showed that 2.2 pN of force is often required for penetrating agar that resembles skin dermis^16^. Our results showed that actual *Leptospira* cells could produce a swimming force around eightfold higher than the previous model predicted. Furthermore, the swimming force of *Leptospira* is ∼30 times as much as that of *E. coli* (∼0.6 pN)^17^.

**Fig. 3.**
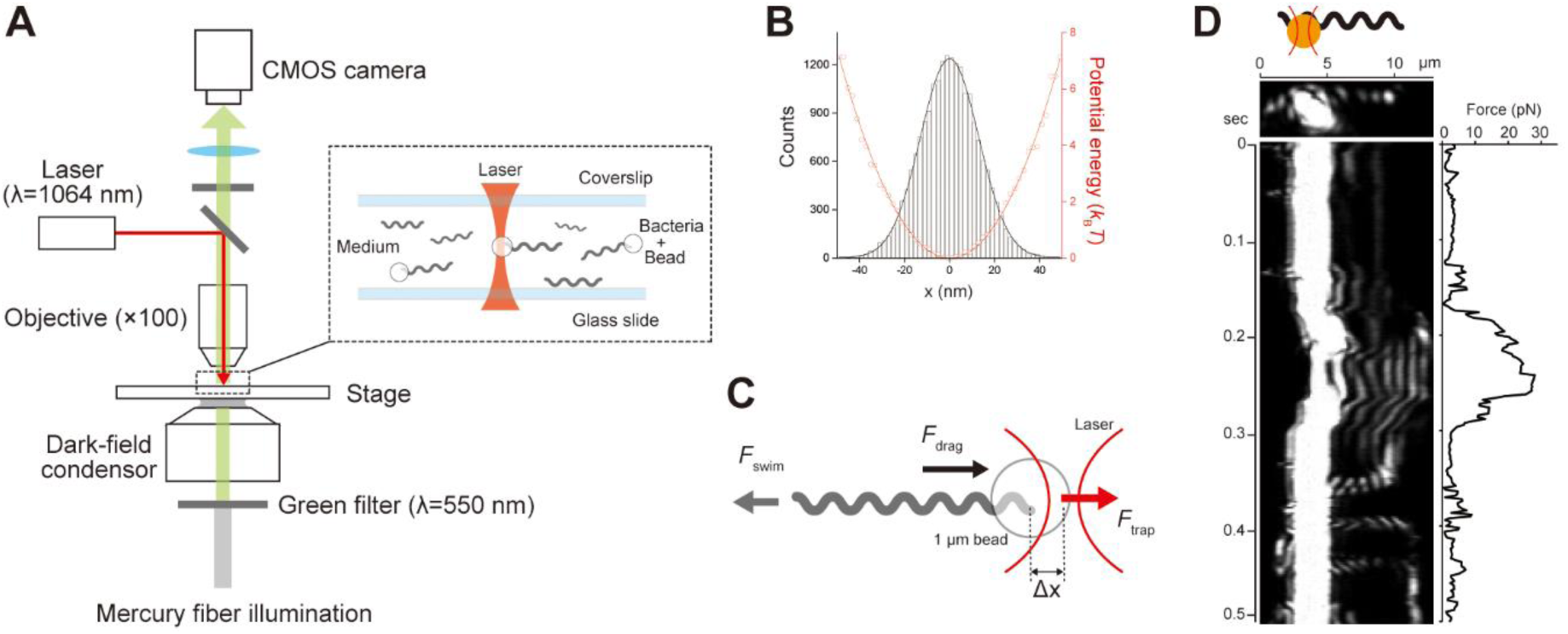
Swimming force measurement. (A) A dark-field microscope equipped with an optical tweezer. (B) The distribution of the positions of a 1-μm bead trapped by optical tweezers (histogram) and the estimated potential profile (red circles). The black and red lines indicate the results of the curve fitting by the Gaussian distribution and harmonic function, respectively. Spring constants were measured in each chamber, and 16-36 pN/μm were used. (C) Force balance in a swimming cell trapped by optical tweezer. See Methods for details. (D) The time course of the bead position is attached to a leptospiral cell and is trapped by an optical tweezer. The left upper schematic represents a trapped *Leptospira* via a 1 μm bead; the left lower panel is a kymograph showing bead movement with the leptospiral swimming. The right panel indicates the time course of the force estimated from the bead movement. See also Video 4.

**Fig. 4.**
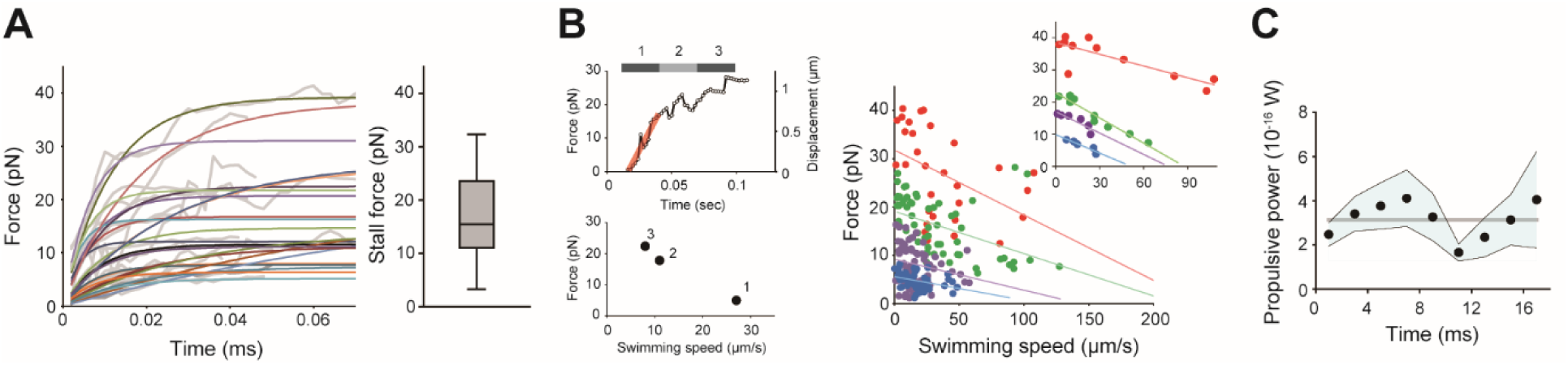
Swimming force. (A) Time courses of the swimming force obtained from 24 *L. kobayashii* cells. Gray and colored lines indicate the measured force curves and the results of exponential curve fitting, respectively. All of the force curves are shown in Supplementary Fig. S1 separately. The right panel shows the stall forces determined by the curve fitting; the boxes show the 25th (the bottom line), 50th (middle), and 75th (top) percentiles, and the vertical bars show the standard deviation. (B, left) Conversion from the force-time plot to the force-speed relationship, which is explained using an example trace extracted from *A*. The data plotted in the force-time plot was separated with an equal time interval, and the averaged forces of each bin were plotted against the average speeds determined by line fitting as shown in red in the upper panel. (right) The force-speed curve obtained from 24 cells were classified into four groups by stall force: <10 pN (blue), 10-20 pN (purple), 20-30 pN (green), and >30 pN (red). The colored lines are the regression lines fitted to each group. Example data extracted from each group are shown in the inset. (C) Time course of the power of the *Leptospira* swimming calculated by *F* · *ν*. Black dots and the blue band indicate average values at each time point (n = 24 cells) and standard error, respectively. The horizontal gray line indicates the average power of the entire time course (3.1×10^−16^ W).

### Force-speed relationship

The relationship between force and speed was obtained from the force-time plots, showing that the swimming forces linearly decreased as the swimming speed increased (Fig. 4B). The force-speed curves suggest that *Leptospira* could vary their propulsive force depending on the load exerted on the cell, such as viscous drag and trapping forces. Namely, the observed variation of the slope in the force-speed curves implies the difference of the load responsivity among cells. We calculated the work per unit time, *F* · *ν*, for the *Leptospira* propulsion using the measured values (Fig. 4C). The propulsive power of *Leptospira* is in the range from 1×10^−16^ to 7×10^−16^ W, which is two orders of magnitude larger than that of *E. coli* (5.8×10^−18^ W)^17^. These results suggest that *Leptospira* can produce a large force while maintaining high efficiency when penetrating gel-like viscous materials.

## Discussion

We focused on the association of motility with the invasion of the *Leptospira* spirochete. Our biophysical experiments revealed that the back-and-forth movement of *Leptospira* is enhanced when only one of the cellular ends is exposed to high viscosity and that the spirochete can produce a much larger swimming force than exoflagellated bacteria. The trial-and-error behavior that gives the cell to find preferable entrances will be significant for infection through the dermis, because such heterogeneous fibrous structures may obstruct the spirochetal invasion even though they can elicit high propulsion forces. The viscosity-dependent swimming reversal was observed in both pathogenic and non-pathogenic species, implying that *Leptospires* can penetrate the dermis regardless of pathogenicity kinematically. However, since the motility of the non-pathogenic *L. biflexa* is lost immediately after exposure to physiological osmotic conditions^18^, dissemination within the host will not be possible.

Enhancement of swimming reversal has been observed in polymer solutions, where the entire spirochetal body was exposed to changes in environmental viscosity or viscoelasticity^12^. However, we showed that swimming reversal is increased by a minimal part of the cell body that is placed in the viscoelastic milieu. Given that the torque for rotating the spiral body of *Leptospira* is produced by PFs residing at both cellular ends, the two PFs should rotate cooperatively. In this context, our results suggest some sort of signal transduction between the two cell ends. Thus, one of the remaining issues to be elucidated is the mechanism by which *Leptospires* sense the viscoelastic change at one end and rapidly transmit the mechanical signal to the other end (<1 s at the latest; see Video 5 for the rapid reversal). It is known that coordinated rotations observed between *E. coli* flagellar motors depend on diffusion of the phosphorylated chemotactic signaling protein, CheY-P^19^. *Leptospira* spp. also possess several CheY homologues^20^. However, the formula *t* = *x*^2^/*D* depicts that for CheY with a diffusion coefficient *D* = 10 μm^2^/s^19^, it is estimated to take ∼40 s (t) for diffusing *x* = 20 μm (approximate distance between the leptospiral motors). Therefore, rather than such cytoplasmic signaling, the relatively stiff protoplasmic cylinder may be a possible medium for mechanical signal transduction^11,21^. Elucidating the coordinated control mechanism of PFs warrants further investigation.

The measured swimming force of *Leptospira* was ∼30 times larger than that of *E. coli*^17^. Such higher force generation by the spirochete could be attributed to drag coefficients and high torque from the motor. Hydrodynamic studies of low Reynolds number using the resistive force theory showed that drag force exerted on a spherical cell body with a spiral, thin body, that rotates at *ω*_Lep_ and translates at *ν*_Lep_ in liquid is calculated by *F*_Lep_ = *α*_Lep_*ν*_Lep_ + *β*_Lep_*ω*_Lep_., where *α*_Lep_ and *β*_Lep_ are drag coefficients for a spiral cell body^6,11,22^. Similarly, the drag force on an externally flagellated bacteria assumed by a spherical body with a flagellum is given by *F*_Efb_ = *α*_Efb_*ν*_Efb_ + *β*_Ef_*ω*_Ef_, where *α*_Efb_ is the sum of the drag coefficients for the translating spherical cell body, *α*_*Cell*_, and filament *α*_Ef_ (*α*_Efb_ = *α*_*Cell*_ + *α*_Ef_), and *β*_Ef_ is the drag coefficient for the flagellar rotation; *ν*_Efb_ and *ω*_Ef_ are the swimming speed and the flagellar rotation rate, respectively^23^. These drag coefficients depend on geometrical parameters such as the length and width of a cell body, the wavelength and the amplitude of a helix and fluid viscosity (see Methods). Calculations using morphological parameters of *L. biflexa*^11^ and *E. coli* give *α*_Lep_ = −0.056 pN·s/μm, *β*_Lep_ = 0.002 pN·s, *α*_Ef_ = −0.02 pN·s/μm, and *β*_Ef_ = 0.0003 pN·s. Thus, these results indicate that drag coefficients for the leptospiral body is larger than those for *E. coli*. Despite such large drag coefficients, the swimming speed (∼20 μm/s) and cell-body rotation rates (>100 Hz) of *Leptospira* spp. are comparable to those of *E. coli*^17^. Cryo-electron microscopy revealed that, in comparison with conventional flagellar motors such as *E. coli* and *Salmonella* spp, the rotor ring of the spirochetal flagellar motor is larger^24^, and that a greater number of torque generators (stator units) are assembled^25^. Thus, the large swimming force of *Leptospira* could be provided by high torque from the motor that can rotate the heavy cell body. Since the cell morphology, the motor structure, and the swimming velocity are very similar among different species of *Leptospira*^18,25^, the spirochetal genus could be characterized as being powerful swimmers.

The stall force was uneven among the measured cells (Fig. 4A). At high load, the motor torque of the external flagellum depends on the number of the stators and the input energy ion motive force (IMF)^26^. Due to the stable incorporation of the full number of the stator units to the motor, shown in *Leptospira* spp. by electron cryo-tomography^25^, the difference in the IMF is a possible cause of the varied stall force. Furthermore, since intimate contact between PFs and cell membranes is thought to be necessary for spirochetal swimming^27^, the PF-membrane interaction or morphological differences of PFs, such as length, could affect the propulsion.

The force-speed relationship (Fig. 4B) showed that the swimming force decreases with increased speed. This result suggests the mechanism by which the spirochete varies the propulsive output in response to the change in load, producing more significant force at high load. The load-dependent assembly of the stator units is observed in the flagellar motors of *E. coli* and *Salmonella*^28–31^. Although such stator dynamics seem to be implausible for the *Leptospira* motor as referred above, *Leptospira* may adjust their propulsive force through another mechano-sensing mechanism, allowing the spirochete to invade highly viscous environments while maintaining large power output.

## Methods

### Bacteria and media

The saprophytes *Leptospira biflexa* strain Patoc I and *Leptospira kobayashii*^32^, and the pathogenic species *Leptospira interrogans* serovar Manilae strain UP-MMC-NIID^33^ were grown at 30°C for 4 days in Ellinghausen-McCullough-Johnson-Harris (EMJH) liquid medium with 10% bovine serum albumin until the stationary phase. Potassium phosphate buffer (20 mM, pH7.4) was used as a motility medium^12^.

### Measurement of swimming reversal

The *Leptospira* culture was centrifuged at 1,000*g* for 10 min at 23°C, and the precipitated cells were resuspended in the motility medium without dilution. The bacterial suspension was infused to a flow chamber that was made by sticking a coverslip and a glass slide with double-sided tape (90 μm in thickness) that contained 0.2% agar so that the agar and liquid area were contiguous in the chamber (Fig. 2A). The liquid-agar border was observed with a dark-field microscope (BX53, 40× objective, 5× relay lens, Olympus, Tokyo, Japan), and behaviors of cells inserting one end of the cell body into agar were recorded with the CMOS camera at 250 Hz.

The swimming reversal was measured by tracing the cellular centroid in general^34,35^. However, the morphology of the *Leptospira* cell changes frequently, thus affecting the consistency between the centroid displacement and the actual cell movement (Supplementary Fig. S2). The positions of both ends of the cell body were determined together with the centroid, and simultaneous displacements of the three points were recognized as cell movements to avoid the false recognition of the reversal (Fig. S2A). Swimming speeds were measured by line fitting to the time courses of the cell displacements at an interval of 0.1 s, <1 μm/s was judged as “pausing”. The reversal frequency was determined by normalizing the number of the reversals (*N*_*reν*_) by the observation time (*t*), such that *N*_*reν*_/*t*. The data were analyzed with ImageJ software (National Institutes of Health, Rockville, MD) and programs originally developed using LabVIEW 2014 (National Instruments).

### Measurement of swimming force

A dark-field microscope (BX50, Olympus, Tokyo, Japan) that is essential for observing a thin leptospiral cell body (∼140 nm in diameter) was equipped with an optical tweezer (Fig. 2A). The cell suspension prepared by the same procedure as the reversal measurement was mixed with 1.0 μm carboxyl latex beads (ThermoFisher Scientific, Waltham, MA) and was incubated at 23°C for 10 minutes. The mixture was infused to the glass-made chamber, and spontaneous bead attachments to swimming cells were observed. The attached bead was trapped by a 1064 nm semiconductor laser (TLD001, Laser Diode Driver, Thorlabs Inc. Newton, NJ) through an ×100 oil immersion objective lens (UPlanFLN, Olympus, Tokyo, Japan), and the bead movement was recorded with a CMOS video camera (acA800-510um, Basler, Ahrensburg, Germany) at a frame rate of 60 Hz. The numerical aperture of the objective was adjusted with the objective-lens aperture to perform dark-field observation and laser trapping simultaneously. The recorded movie was analyzed to determine the bead displacement with a custom-made program developed using LabVIEW 2014 (National Instruments, Austin, TX).

Displacement of a trapped bead (Δ*x*) can be calibrated to a restoring force of optical tweezer (*F*_*trap*_) using the equation *F*_*trap*_ = *k*Δ*x*, where *k* is a spring constant. The values of *k* were determined in each flow chamber by trapping a bead free from cells and analyzing its positional fluctuation. The positional distribution of the trapped bead showed a Gaussian distribution *f*(*x*) ∝ *exp*(–*x*^2^/2*σ*^2^), where *σ* is the standard deviation (Fig. 2B, black line), obeying Boltzmann’s law *P*(*x*) ∝ *exp*(–Δ*U*/*k*_*B*_*T*), where Δ*U* is the potential energy, *k*_*B*_ is the Boltzmann constant (1.38 ×10^−23^ J/K), and *T* is the absolute temperature (296 K), namely, Δ*U* = (*k*_*B*_*T*/2*σ*^2^)*x*^2^ (Fig. 2B, red line). Since the thermal fluctuation of the bead captured by a spring with *k* can be described by the harmonic function *U*(*x*) = 1/2 *kx*^2^, *k* = *k*_*B*_*T*/*σ*^2^. The swimming force was determined using *F*_*trap*_ and *F*_*drag*_ (Fig. 2C). When the bead with a diameter of *r* is moved at a speed of *ν* (swimming speed of the cell) in a solution with a viscosity of *μ, F*_*drag*_ = 6*πμr* · *ν*, where 6*πμr* is a drag coefficient given by Stokes’ low. The viscosity of the motility medium was measured with a tuning-fork-type viscometer (SV-1A, A&D, Tokyo, Japan), giving 0.8 mPa×s at 23°C.

### Drag coefficients

Drag coefficients for the spirochete and externally flagellated bacterium are calculated as described previsouly^11,23^. For the *Leptospira* cell, the protoplasmic cylinder lacking bending at the cell ends was assumed:

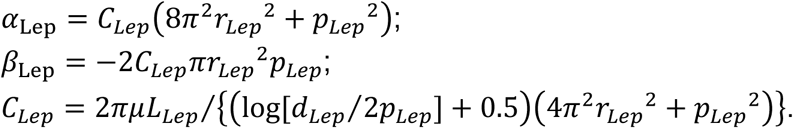

Here, *r*_*Lep*_, *p*_*Lep*_, *L*_*Lep*_, and 2*d*_*Lep*_ are the helix radius (0.09 μm), helix pitch (0.7 μm), length (20 μm), and diameter (0.14 μm) of the leptospiral cell body^11,36^. Drag coefficients for the externally flagellated bacterium that was assumed to consist of a spherical body and a helical filament, referring to the morphology of *Vibrio alginolyticus*^23^, were calculated as follows:

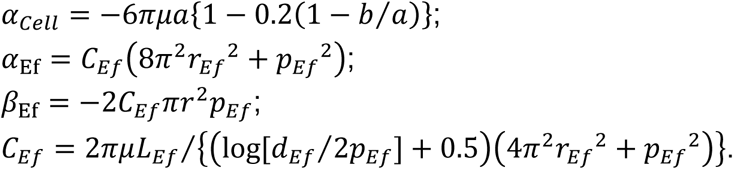

Here, 2*a* and 2*b* are the diameter (0.8 μm) and length (1.92 μm) of the cell body, and *r*_*Ef*_, *p*_*Ef*_, *L*_*Ef*_, and 2*d*_*Ef*_ are the helix radius (0.14 μm), helix pitch (1.58 μm), length (5.02 μm), and diameter (0.032 μm) of the flagellum. The medium viscosity *μ* was assumed to be 1 mPa×s.

## Acknowledgements

We thank Toshiyuki Masuzawa for providing *Leptospira* strains. We also thank Shoichi Toyabe and Jun Xu for technical supports. This work was supported by the JSPS KAKENHI (18K07100 for SN).

## Author contributions

K.A., K.T. and S.N. planned the project; K.A., T.K., K.T. and S.N. carried out the experiments; K.A. and K.T. set up the optical system; K.A. and T.K. made programs for data analysis; K.A., T.K. and S.N. analyzed the data; K.A., T.K. and S.N. wrote the paper; all authors reviewed the manuscript.

## Data availability

The data supporting the findings of this study are available from the corresponding author upon request.

