## Supplementary Information for "Implications of back-and-forth motion and powerful propulsion for spirochetal invasion"

· Supplementary Fig. S1. Swimming force vs time plots.
· Supplementary Fig. S2. Swimming reversal measurement.
· Supplementary Video 1. Trial-and-error and invasion of *L. kobayashii* at the liquid-agar border
· Supplementary Video 2. Swimming reversal of *L. interrogans* at the liquid-agar border
· Supplementary Video 3. Swimming reversal of *L. biflexa* at the liquid-agar border
· Supplementary Video 4. Laser trap of 1-μm polystyrene beads for force measurement
· Supplementary Video 5. Quick swimming reversal of *Leptospira* (slow-motion movie)

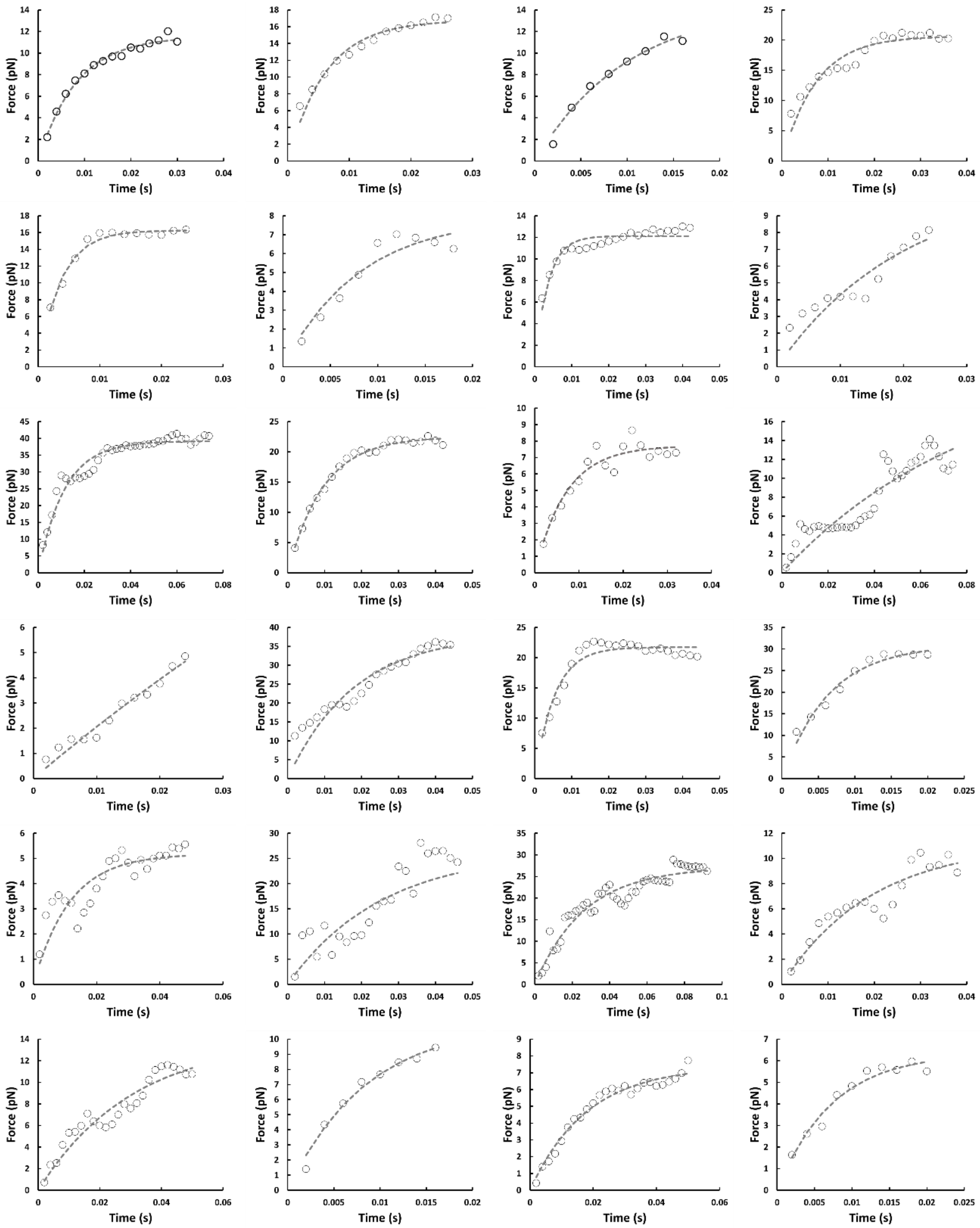

**Supplementary Fig. S1. Swimming force vs time plots.** Circles are experimental data plot, and dashed lines are the results of exponential curve fitting.

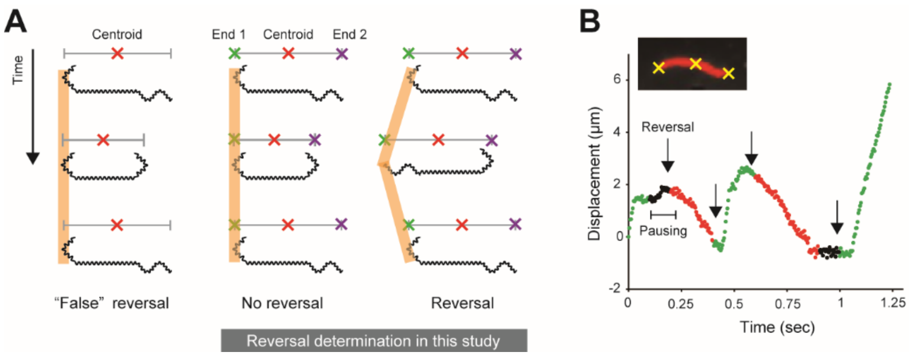


**Supplementary Fig. S2. Swimming reversal measurement.** (A) Determination of swimming reversal. The left panel shows a conventional measurement where the cellular centroids are traced; the center and right panels explain the current method, in which the positions of both cellular ends are determined together with the centroid. Thick orange lines indicate the actual displacement. See Materials and Methods for detailed explanation. (B) Example data of the leptospiral displacement. Green, red, and black indicate the forward movement, backward movement, and pausing, respectively. This data shows four reversals (black arrows). The inset shows the analyzed microscopic image of a leptospiral cell.
